# Curved geometry shapes the observability of hippocampal oscillations

**DOI:** 10.64898/2026.11.05.764490

**Authors:** Julia Makarova, Ricardo Muñoz-Arnaiz, Faezeh Ashouri, Oscar Herreras

## Abstract

Curved neural architectures reshape the spatial distribution of field oscillations, complicating the identification of their underlying generators. To test these principles experimentally, here we resolve the paradoxical poor expression of theta oscillations in the hilus of the folded dentate gyrus (DG), a region that otherwise exhibits some of the largest FPs in the brain. Using laminar recordings across both hippocampi and computational modeling, we show that theta is locally masked by cancellation of phase-coherent contributions. Theta generators in CA1 and DG produce volume-conducted fields that, due to the curved geometry of the DG, exhibit opposite dipolar orientations in the hilus, leading to regional cancellation. In contrast, beta and gamma generators are only weakly coordinated across hippocampal regions and therefore remain prominently expressed despite the same geometry. These results demonstrate that theta amplitude, local observability, spatial segregation and apparent translocation emerge from the interplay between geometry and temporal coherence among distributed generators, with tissue geometry acting as a spatial filter for coherent activity.

**Significance Statement:** Brain potentials are widely used to infer neuronal activity, yet how source geometry shapes their spatial expression remains poorly understood. We show that curved brain structures can reshape where oscillations appear to originate. In the dentate gyrus hilus, theta activity is not truly absent but masked by geometry-dependent cancellation between coherent sources in CA1 and DG. In contrast, the same geometry amplifies non-coherent beta and gamma oscillations, making them prominently visible. These findings demonstrate that the apparent location of oscillations can diverge from their true sources, revealing that geometry and temporal coherence jointly shape what is observable in brain potentials.

## Introduction

Field potentials (FPs) are widely used as neuronal correlates of population activity during behavior and in pathological states. The formation of extracellular fields reflects interactions operating at multiple spatial scales: extensive cancellation within individual generators and superposition among distributed generators, both of which depend on the spatiotemporal organization of the underlying currents (Lorente de Nó, 1947). Consequently, the amplitude, polarity, and even the regional detectability of FPs may be strongly shaped by the combined three-dimensional geometry of activated neuronal populations (Nunez and Srinivasan 2006; Martín-Vázquez et al., 2016; Herreras et al., 2023a). Despite these theoretical principles, empirical demonstrations of how geometry shapes the interpretation of extracellular recordings remain scarce.

The hippocampus is a convenient model system because its laminar organization combines well-defined current generators with a highly folded three-dimensional architecture. This is particularly relevant because tissue curvature can displace the center of mass of FP expression away from the underlying generators, potentially leading to source mislocalization and unexpected mixtures of signals at a given recording site (Woodbury, 1960; Fernández-Ruiz et al., 2013; Gabay and Robinson, 2017; Torres et al., 2019; Herreras et al., 2023a). The dentate hilus offers a particularly informative test case because it lacks the cytoarchitectural organization required for the emergence of mesoscopic field generators (Lorente de Nó, 1947), yet it exhibits some of the largest FPs recorded in the brain (Fernández-Ruiz et al., 2013).

Hippocampal oscillations provide a suitable probe because they arise from multiple generators distributed throughout the hippocampal formation, and have sufficient spatial extent to allow geometry-dependent distortions be experimentally resolved (Benito et al., 2014). Theta oscillations (4–8 Hz) are present across CA1 and DG and involve several spatially distributed generators that become highly synchronized through common septal drive (Leung, 1984b; Brankack et al., 1993; Kocsis et al., 1999; Montgomery et al., 2009; López-Madrona et al., 2020). Other hippocampal oscillations, including beta and gamma rhythms, provide complementary situations in which distributed generators may differ in their spatial extent or the degree of coherence and therefore in their extracellular expression (Lockmann et al., 2018; Csicsvari et al., 2003; Benito et al., 2016). The circuit mechanisms underlying these oscillations have been extensively studied, although their interpretation remains complicated by the contribution of multiple generators and interneuron populations (Sinclair et al., 1982; O’Keefe and Reece, 1983; Fox et al., 1986; Vertes and Kocsis, 1997; Csicsvari et al., 1999; Buzsáki, 2002; Czurko et al., 2011; Colgin, 2013; Huh et al., 2016; Korotkova et al., 2018). While these mechanisms are essential for understanding how oscillatory activity emerges, the central question addressed here is not how oscillations are generated, but how they become represented in extracellular recordings.

Experimental theta profiles show maxima at locations that do not necessarily correspond to regions of strongest synaptic input, whereas other areas with substantial afferent drive (e.g., CA3) may exhibit weak or negligible theta FP expression (Tóth et al., 1997). A striking example is the poor expression of theta oscillations in the hilus of the dentate gyrus (DG) under anesthesia (Bland and Wishaw, 1976). This observation is paradoxical because the hilus is embedded within the folded dentate architecture and it displays large beta and gamma oscillations and other fast activity patterns.

Using laminar recordings across both hippocampi together with computational modeling, we investigate how the curved geometry of the DG shapes the extracellular expression of theta and other hippocampal oscillations. We show that interactions between source geometry and temporal coherence among coactivated generators can mask or enhance oscillatory signals and produce spatial patterns that do not directly reflect the location of their underlying sources.

## Results

Here we use the term source (of current) to refer to an anatomical region that produces coherent potentials, i.e. FP-generators. To establish a reference map of hippocampal field generators, we first characterized FP organization during irregular activity (this analysis is provided in Supplementary Fig. S1; see also Herreras et al., 2015, Hernández-Recio et al., 2023, and references therein). Under this condition, current sources and sinks appeared as spatially discrete events that could be more readily assigned to anatomical layers using current source density analysis (CSD; Nicholson and Freeman, 1975), whereas independent component analysis (ICA; Makarov et al., 2010) of laminar FPs facilitated the separation of spatially coherent components (FP-generators) and the characterization of their spatial extent and gradients (Benito et al., 2014).

### Synchronous activation across DG blades renders giant hilar potentials for beta but not theta currents

Whereas synchronous activation of opposing granule cell dipoles across the DG blades can generate giant hilar potentials during non-theta states (Fernández-Ruiz et al., 2013), this phenomenon was paradoxically absent during theta (Fig. 1A). To address this discrepancy, we examined the laminar organization and interregional coherence of theta-, beta-, and gamma-associated FPs and their underlying currents and ICA-derived FP generators during theta states.

**Figure 1.**
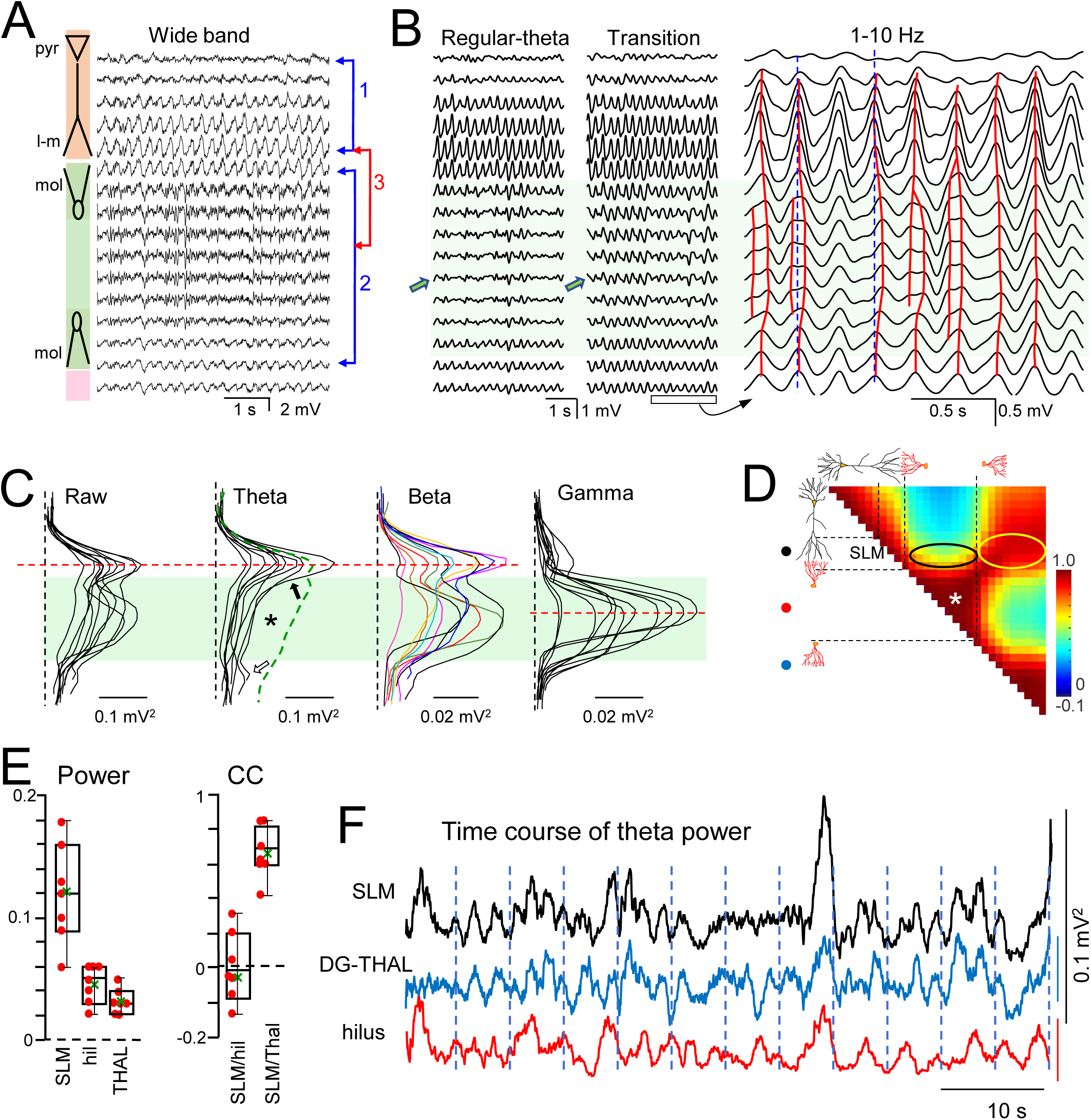
Spatial and temporal features of FPs during theta states. A, Sample epoch. Prominent theta is observed on both sides of the hippocampal fissure (hf) and in the lower DG blade, whereas the hilus is dominated by faster activity (beta and gamma). Colored arrows indicate sites selected for pairwise statistical comparisons. B, During transitions from irregular to theta states, hilar theta can be observed, albeit with reduced amplitude and spatially inhomogeneous patterns that differ from those at more prominent sites. The 2 s enlargement (right) shows that hilar theta waves follow variable spatial trajectories, in contrast to those in the SLM and lower DG blade/thalamus, which are phase-locked. C, Superposition of mean spatial power profiles for raw FPs and band-pass-filtered signals in the left and right hemispheres (n = 6 animals). For reference, the L–M (theta) ICA component is included (green dashed profile). The asterisk marks the reduction in theta power in the hilus. The black arrow indicates a sharp drop in theta power within the molecular layer of the upper DG blade, followed by a small recovery in the lower blade (open arrow). The dashed red line marks the spatial reference used to align profiles, either at the SLM or at the hilar maximum. Note the frequency-specific spatial profiles. D, Spatial maps of pairwise cross-correlations during regular theta states. Ovals highlight regions of high or reduced correlation. The high internal hilar correlation (white asterisk) is driven by beta and gamma activity. E, Group statistics for mean power and cross-correlation at selected sites. F, Temporal envelopes of theta power at three sites (theta-filtered FPs).

Spontaneous and sensory-evoked theta epochs exhibited similar features and were analyzed together, with quantification restricted to the left hippocampus. Mean theta frequency was 4.03 ± 0.17 Hz (n = 7) and did not correlate with the respiratory rhythm (1–2 Hz; Ylinen et al., 1995; Lockmann et al., 2016).

During transitional epochs from irregular to theta states, short intermittent theta bouts emerged across the hippocampal fissure, from the CA1 stratum lacunosum-moleculare (SLM) to the upper molecular layer of the DG (Fig. 1B). These brief, unstable transitions constituted an exception to the otherwise negligible hilar theta expression observed under anesthesia. In the raw traces, hilar theta remained barely visible, but theta-band filtering uncovered a substantial oscillatory component, indicating that it had been partially obscured by concurrent large-amplitude beta activity. Successive theta waves in these episodes varied in phase across the hilus, and some appeared to bifurcate from the upper DG blade before converging or fading near the lower DG blade (Fig. 1B). The cycle-to-cycle phase variability across the hilus is difficult to reconcile with a single remote generator and instead suggests changing contributions from multiple incompletely phase-locked generators.

In epochs of stable regular theta, beta and gamma but not theta oscillations were present in the hilus; we then explored these bands comparatively, starting with their spatial coverage. Band-pass filtered FPs were used to derive spatial profiles of mean power across frequency bands (power profiles in Fig. 1C; left and right hippocampus, n = 6 animals). Along the CA1–DG linear track, theta power peaked near the hippocampal fissure (red dashed line), and declined sharply within the upper DG blade (black arrow). Power was markedly reduced in the hilus (asterisk), where it continued to decrease slightly, before modestly re-emerging in the lower DG blade (empty arrow). From this point ventrally, theta power resumed its downward trend as recordings extended into the thalamus (Fig. 1A). Gamma power was maximal in the hilus, with a smaller and only occasionally detectable secondary peak in CA1 SLM.

In contrast, beta-band activity displayed two well discriminated peaks of power, one in CA1 SLM and another in the DG hilus, with highly variable relative amplitude. Although a weak inverse trend was detected (linear regression slope = −0.2), CA1 and DG beta amplitudes were not significantly correlated (r = 0.28). Neither beta nor gamma power profiles showed partial recovery at the lower DG blade that might suggest hilar discontinuity.

Spatial coherence between nearby and distant regions was estimated through a spatial matrix of pairwise correlations between recording sites (Pearson’s linear coefficients: Fig. 1D). Spatial correlation maps of wideband FPs during theta were similar to those observed in irregular states (see SF-1). The hilus exhibited high intrinsic correlation (Fig. 1D, asterisk), whereas its coupling with SLM was markedly reduced (black oval), suggesting that is primarily driven by non-theta activity (e.g., beta-dominant signals). In contrast, correlation between CA1 SLM and sites beyond the hilus (lower DG blade and thalamus) approached maximal values (yellow oval), consistent with intrinsic coherence.

We compared theta-filtered FPs across three sites (SLM, hilus, and DG–thalamus border; Fig. 1E). One-way ANOVA revealed significant differences in mean power (mean ± SEM: 0.12 ± 0.015, 0.044 ± 0.005, 0.031 ± 0.004; n = 7; F(2, 18) = 23.55, p = 9.46 × 10 ⁶), with Tukey post hoc differences revealing significant difference between SLM and hilus (p = 3.3 × 10 ⁴) and between SLM and DG–thalamus border (p = 1.8 × 10 ⁵), but not between the latter two groups (p = 0.6339). These values are consistent with a spatial decay from SLM; however, correlation analyses revealed an additional functional structure. SLM–thalamus coupling was strong (r = 0.67 ± 0.05), whereas SLM–hilus coupling was weak and variable (r = 0.02 ± 0.08). Group differences were assessed using a paired t-test on Fisher z-transformed correlation coefficients, revealing a significant difference (t(6) = 10.7, p ≈ 2 × 10 ⁵).

This differential hilar involvement was also reflected in the temporal fluctuations of mean theta power (Δt = 5 s), which were highly similar between SLM and thalamic sites but more independent in the hilus (Fig. 1F), suggesting a distinct contribution of local theta-related processes in this region.

### CA1 and DG theta generators are coherent; beta/gamma are not

We further explored ICA-derived FP generators, as this approach provides a spatial decomposition of extracellular activity into independent components associated with coherent population processes (Herreras et al., 2015). The canonical spatial profiles of the main hippocampal FP generators identified during irregular activity are shown in Fig. 2A (see also Supplementary Fig. S1), together with the corresponding CSD profiles from a representative epoch. These reference profiles provide a stable framework across animals to compare generator organization during theta states (Fig. 2B). Several of these components have been associated with pathway-specific activities, including the CA3-to-CA1 Schaffer generator and the medial and lateral entorhinal inputs to the granule cell population (MPP and LPP generators, respectively) (Benito et al., 2014). During theta, activity was dominated by a distinct CA1 generator with a maximum in the SLM (LM generator, green traces, Fig. 2C). A weaker theta modulation was also detected in the CA1 Schaffer generator in five of seven experiments. In the DG, theta was consistently captured by a single generator (cyan traces, Fig. 2C), which in four of seven experiments displayed a spatial profile compatible with the MPP generator. This component showed greater spatial variability, consistent with the smaller contribution of this particular component to the variance structure used for ICA extraction (Herreras et al., 2015). These results indicate that hilar theta suppression cannot be attributed to the absence of a DG theta-related generator, but rather to the way distributed generators combine in the extracellular space.

**Figure 2.**
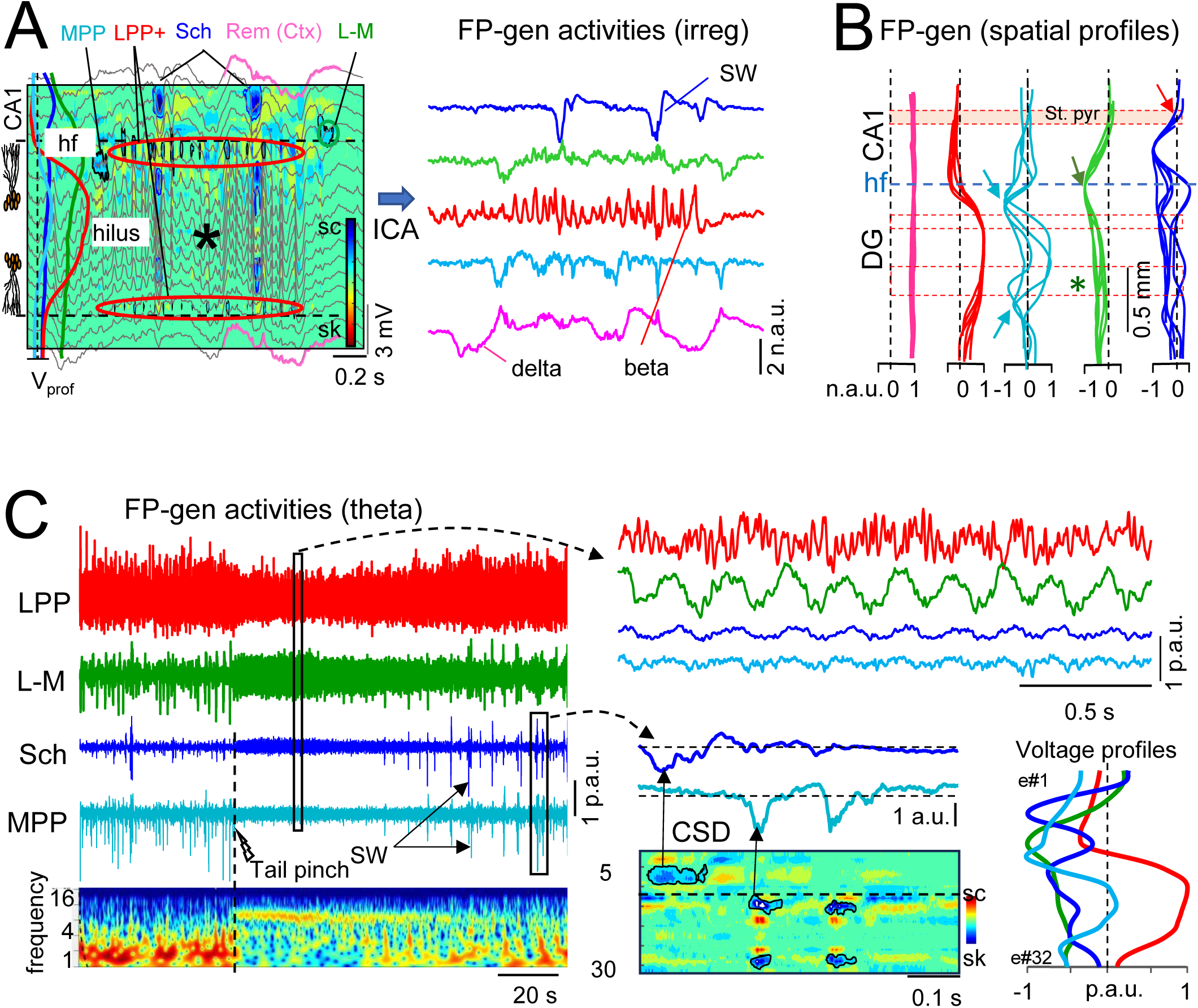
Activity of FP generators disentangled by Independent Component Analysis (ICA) during irregular and theta states. A, Upper panel: superposition of raw FPs, CSD, and voltage profiles (Vprof) from FP generators (left), across the hilus during a 2 s epoch of irregular activity with strong volume-conducted cortical waves throughout the DG (pink traces). Note the absence of CSD transients in the hilus despite the presence of large FPs (asterisk). Most of these correspond to a prototypical sequence of beta waves, with currents observed in GC domains above and below (red ovals), where potentials have already dropped markedly. Isolated current transients appear at specific strata and coincide temporally with fluctuations in ICA components (right panel), displaying well-defined component dynamics. p.u., proportional units. B, Superposition of spatial profiles of ICA components from six animals (four for the MPP generator), whose stability indicates their structural basis. Profiles were aligned at the CA1 border using the LM generator peak (SLM; green arrow). The more variable MPP generator often exhibited two local minima in the molecular layers (cyan arrows). Profiles are not anatomically re-scaled, and horizontal red dashed lines mark the limits of cell layers across experiments. Gcl, granule cell layer. C, Main and secondary theta generators in ICA components. The ICA analysis corresponds to a representative epoch showing a transition to the theta state (tail pinch) and subsequent recovery to the irregular state (see spectrogram below for the FP recorded at the SLM). During theta, the activity of individual FP generators changes in distinct ways. The dominant irregular activity in the LPP generator is moderately reduced, while still exhibiting strong beta oscillations. In contrast, the L–M generator becomes highly regular and dominant, displaying prominent theta and gamma activities. In this experiment, weaker theta activity is also observed in the Schaffer and MPP generators, whose associated sharp waves disappear. Expanded segments of theta and irregular patterns are shown on the right, together with a CSD contour plot to localize source–sink currents. Spatial profiles computed over a 60 s epoch (right) demonstrate the stability of these activity patterns and support their identification.

The macroscopic spatial features of currents for theta and beta are depicted for a sample epoch at different time scales in figures 3A and 3B, respectively. As synchronicity is a major factor defining the outcome of signals mixing in the volume, we examined phase relationships using the phase-locked value (PLV) parameter (significance was assessed using a surrogate test, n = 1000; p < 0.05) and the CC function. The CA1 exhibited two prominent out of phase theta current dipoles (sites marked as *1* in Fig. 1A): PLV = 0.84 ± 0.01, (phase lag = 2.05 ± 0.18 radians, i.e. ≈ 81 ms; n=5), and CC = 0.42 ± 0.02 (Tmax = 103.6 ± 23 ms). In the DG, two others theta dipoles were found, one in each DG blade, which spanned across all GC domains. For estimation of synchrony between sites we used either FPs, CSD transients, or both. For DG theta FPs (sites marked as *2* in Fig. 1A) the PLV was 0.99 ± 0.01 (phase lag = 0.04 ± 0.01 rad, i.e. ≈ 0.4 ms), and for theta currents (estimated at the outer third in both blades) it was 0.78 ± 0.02 (phase lag = 0.12 ± 0.08 rad, i.e. ≈ 5 ms). That is, theta activity is highly synchronous across DG blades; given the inverted dipole configuration in both blades with currents of the same polarity facing the hilus, this should have promoted large hilar theta potentials.

**Figure 3.**
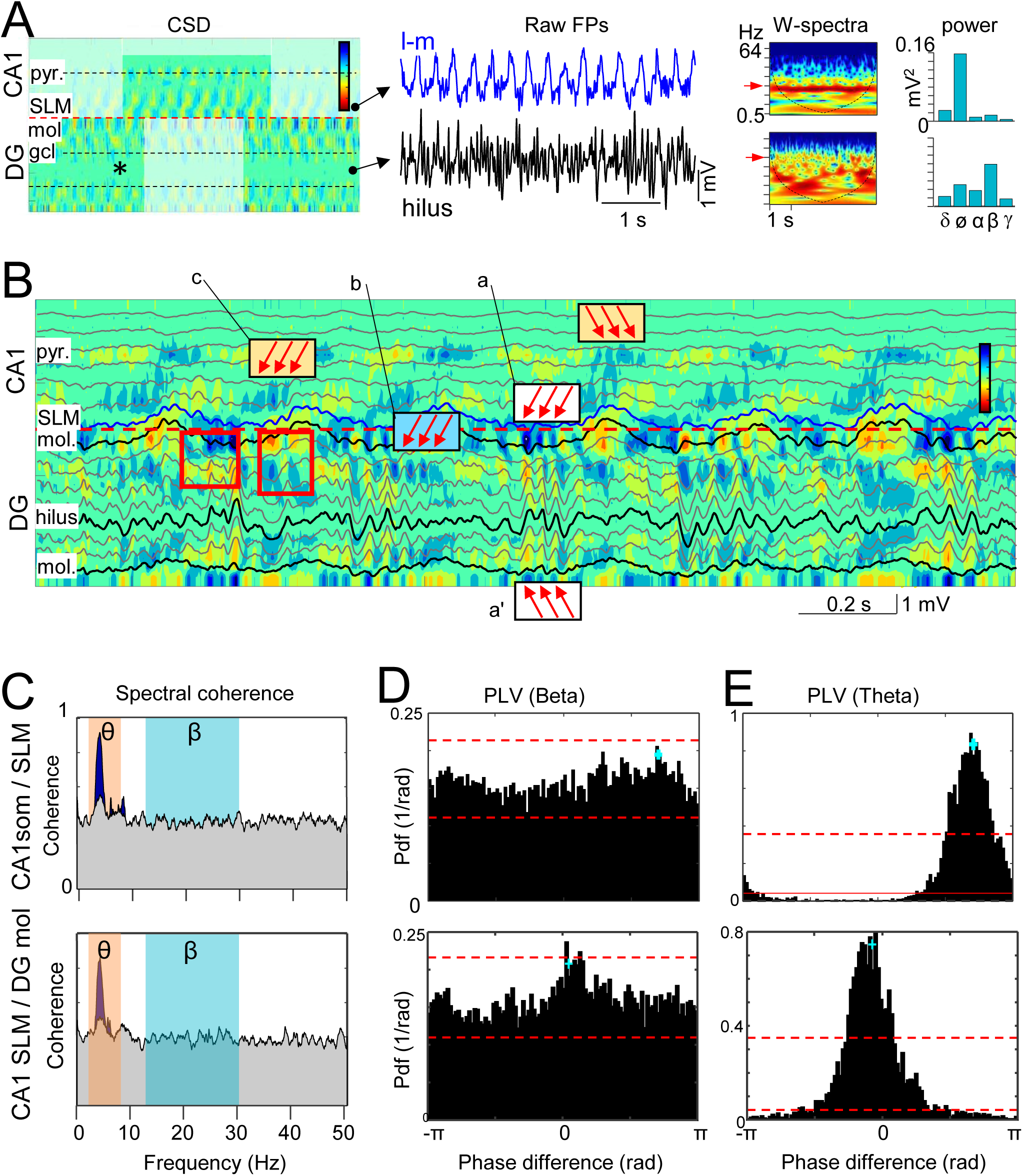
Spatial distribution and interregional coherence of theta and beta currents. A, CSD analysis over a 5 s epoch across CA1 and DG. The red line marks the hippocampal fissure, and black lines indicate cell body layers in both regions. Shaded zones facilitate visualization of each region. In CA1, two pairs of theta source–sink dipoles are observed at the apical distal dendrites and cell body layers, with a strong phase lag between them. In the DG, two sets of theta dipoles are present, one in each blade. Alternating source–sink patterns prevent reliable determination of polarity. No significant currents are detected in the hilus (asterisk), despite large raw FPs (examples shown to the right), indicating volume conduction. Spectral content at each location is illustrated in wavelet spectra and quantified using periodograms. B, Enlarged CSD plots allow visualization of theta and beta activity across regions and layers. Colored boxes (three-arrow markers) highlight trains of beta sinks in different strata of both regions, typically riding on theta current envelopes (red square). However, only the DG hilus displays large beta FPs (superimposed traces). C, Spectral coherence of currents between selected pairs of sites marked in Fig. 3A, within CA1 (st. pyr and SLM, upper row) and across regions (CA1 SLM and DGmol, lower row). Blue peaks above the gray band indicate significant coupling at theta frequencies (surrogate test, n=1000). D, E, Phase-locking value (PLV) for beta (D) and theta (E) currents for the same site pairs. Red dashed lines indicate significance thresholds (surrogate test, n = 1000).

In turn, beta CSD transients exhibited a similar dipolar configuration and orientation as theta CSD transients in the DG, although they were restricted to the middle and outer thirds of the molecular layer and were less synchronous across blades than theta transients (CSDs: PLV = 0.67 ± 0.01; phase lag = 0.27 ± 0.12 rad, i.e., ≈2 ms; n = 5). Beta currents were also observed in CA1 and were primarily centered in st. rad and pyr (Fig. 3B, arrows labeled c), occasionally also in SLM, but similarly to somatic CA1 theta currents they were not associated with significant FPs.

Next, we analyzed these activities along the CA1–DG axis, focusing on CA1 SLM and the outer third of the DG molecular layer, where the main theta dipoles reside. To avoid confounds from tissue curvature, we included only experiments in which CA1 and DG currents were clearly separable during irregular activity. The CA1–DG border was functionally defined by the convergence of: (1) a discontinuity in CSD transients reflecting distinct generators (CA1 distal dendrites vs DG granule cells; Fig. SF1), (2) the peak of the ICA LM generator at the apical CA1 dendrites, and (3) the outer rim of the CSD transient evoked by LOT stimulation in the upper DG blade (LPP input: Fig. SF1).

We computed spectral coherence on CSD signals to assess frequency-specific linear coupling (Fig. 3C). Significant coherence was confined to the theta band, whereas alpha, beta, and gamma-band currents did not exhibit significant coherence between CA1 (SLM or st. pyr) and DG molecular layer (surrogate test, n = 1000). These results were repeated in all five animals tested. We then estimated the PLV amongst pairs of sites. Between CA1 SLM and DGmol sites, the PLV for theta currents was 0.78 ± 0.02 (phase lag = 0.25 ± 0.05 rad; ≈10 ms), whereas for beta currents it dropped to 0.12 ± 0.03 (n = 5; none exceeded significance in the surrogate test; phase lag = 0.33 ± 0.09 rad; ≈2.6 ms; Fig. 3D). We also estimated the CC that was significant for theta currents between CA1 SLM and DGmol (0.44 ± 0.01; Tmax = 3.25 ± 2.14 ms) but not beta currents. In addition, the probability density functions showed much higher percentage of samples above significance level for theta than for beta pairs (Figs. 3D, E), indicating that interregional synchronization of beta waves was less persistent than for theta waves. These findings establish differential cross-regional phase synchronization and persistence in theta but not beta waves. Thus, although activity in other frequency bands is generated both in CA1 and DG, only theta showed interregional coupling.

To assess coordination between DG and CA1 gamma while minimizing volume-conduction effects, phase locking was quantified between the ICA-derived LM and LPP generators, peaking at both sites of the hippocampal fissure. Mean PLV was low, although significantly above surrogate values in all animals (0.176 ± 0.040 for low gamma and 0.135 ± 0.033 for high gamma). In contrast, neither band exhibited significant spectral coherence. These findings indicate weak and transient coordination between gamma generators, in marked contrast to the coherent organization of theta generators.

### Functional blockade of entorhinal inputs reveals the contribution of volume-conducted CA1 theta in the DG hilus

The above results suggest that a theta-paced generator in the DG may counteract the passive decay of theta potentials arising in CA1 SLM as they propagate passively through the DG. To test this, we blocked direct entorhinal inputs to the DG by injecting small volumes of lidocaine into the angular bundle (AB) in one hemisphere while recording from both. In five of seven experiments, complete blockade of the AB-evoked potential was achieved in the ipsilateral hemisphere. Spontaneous activity changes remained largely restricted to the ipsilateral site (Fig. 4). The main observation was that lidocaine unmasked theta activity in the hilus ipsilateral to the injection, whereas beta activity was markedly reduced. In contrast, the contralateral hilus remained unaffected (Fig. 4A).

**Figure 4.**
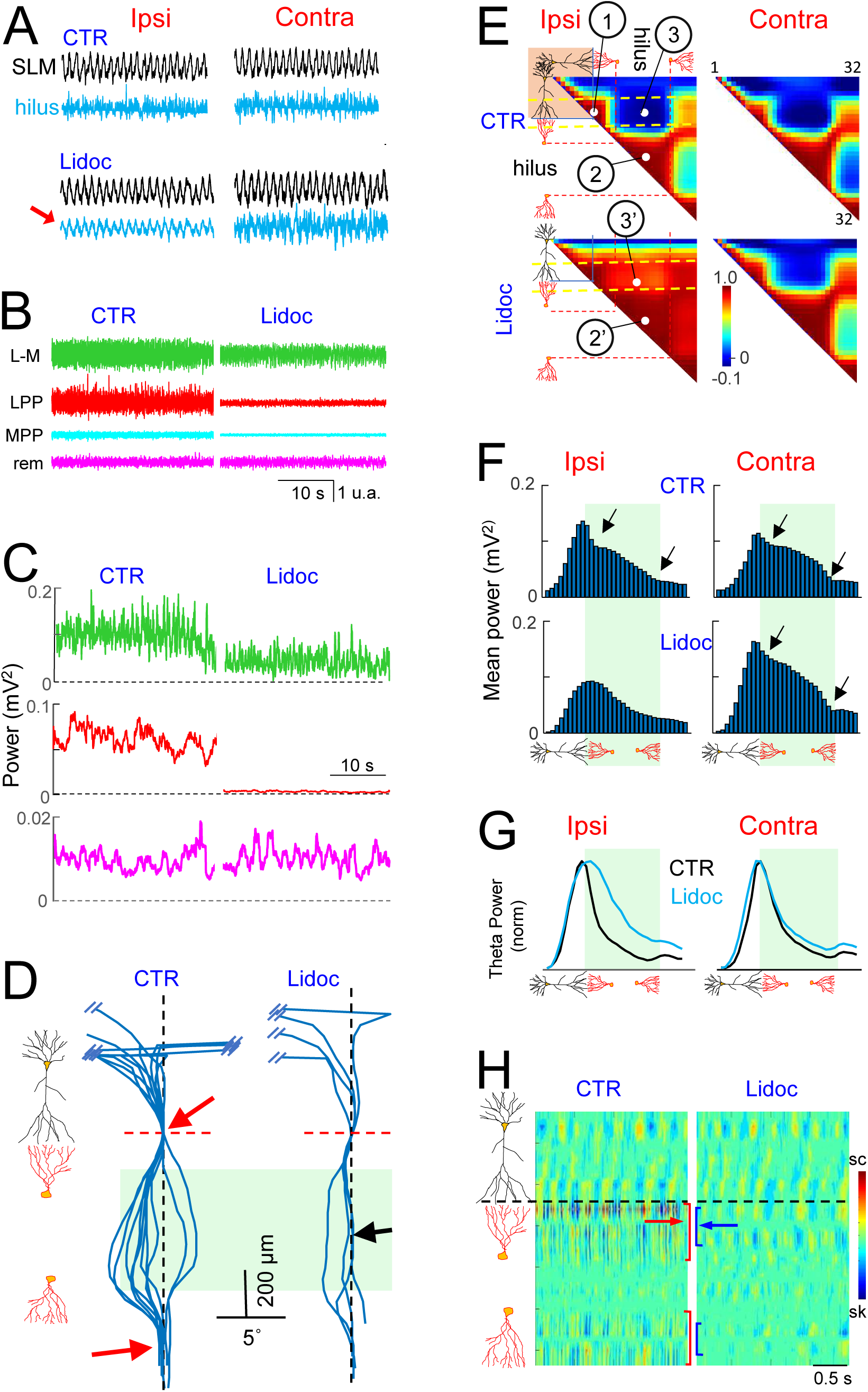
Removing DG input reveals hidden CA1 theta. A, Sample FP traces from the SLM and hilus in both hemispheres before and after lidocaine injection into the left AB. Note the ipsilateral unmasking of theta (red arrow). B, Representative epoch of FP generators before and after lidocaine. The extended time scale facilitates visualization of the selective blockade of entorhinal-driven generators (LPP and MPP), whereas the remote (cortical) generator remains unaffected. C, Temporal envelope of power for selected FP generators ipsilateral to the entorhinal input blockade. D, Phase coherence of theta waves in theta-filtered FPs along a vertical track across multiple experiments, before and after entorhinal blockade. Plots are aligned at the SLM, which serves as reference (dashed line). Some tracks are truncated, and the width of the green hilar shading is approximate. Note the tight phase-locking between the SLM and deeper DG sites under control conditions (red arrows), which also affects the hilus after entorhinal blockade (black arrow). Control: n = 5 (left and right); lidocaine: n = 4 (ipsilateral only). E, Spatial maps of pairwise FP correlations show widespread homogenization after entorhinal blockade, as theta becomes unmasked in the hilus and correlates across sites . F, Selective changes in the theta power profile in the hippocampus ipsilateral to entorhinal blockade. The disappearance of local minima (arrows) in the DG molecular layers results in a unimodal profile. G, Normalized theta power profiles show increased hilar power relative to the SLM peak after entorhinal blockade, consistent with hilar theta unmasking. H, Representative CSD contour plots before and after lidocaine. In this experiment, residual theta currents are still observed in the DG, showing a narrower spatial distribution (brackets) and a shift in the reversal site (arrows). Additional experiments are shown in SF2.

During theta states, the following effects (Fig. 4) were consistently observed across multiple measures in all animals (see additional experiments in Fig. SF2): (a) a reduction in the activity of specific FP generators, pronounced in the DG (LPP, MPP) and modest in CA1 SLM (Figs. 4B,C); (b) a reduction in the phase difference between hilar and SLM theta potentials (reliable phase estimates could only be obtained in n = 4 experiments; Fig. 4D), which was already small under control conditions (<5°); (c) a marked increase in the CC between raw FPs in the DG hilus and CA1 SLM (zones 3 and 3′ in the spatial map of Pearson correlations; Fig. 4E), reflecting increased theta coherence after lidocaine, whereas beta dominance in the hilus accounted for the lack of correlation under control conditions; (d) a shift in the mean spatial power profile across CA1–DG from bimodal to monomodal, suggesting fewer contributing sources, likely one (Fig. 4F); (e) a pronounced widening of the normalized theta power profile toward the hilus, with no change on the CA1 side (Fig. 4G); and (f) a marked reduction in theta and beta currents in the DG (Fig. 4H). This reduction left CSD remnants near the noise level, precluding reliable quantification. Notably, the residual DG theta currents remained approximately phase-locked to those in CA1 SLM. In two experiments, the source–sink dipole appeared shortened, occupying middle and outer thirds of GC dendrites, accompanied by a shift in the reversal point. This may support the presence of at least two DG theta generators under control conditions.

Together, these findings indicate that lidocaine in the AB suppressed DG activity, thereby unmasking the contribution of volume-conducted CA1 signals in the hilus.

### Dipole simulations support DG theta generator masking volume-conducted CA1 theta selectively in the hilus

Because the DG contains multiple generators linked to distinct synaptic pathways, and the sources driving theta currents remain uncertain, we used numerical and FEM models to simulate the mixing of potentials from realistic and hypothetical dipolar configurations to explore their interactions in the volume. For these computations we assumed fully coherent sources to allow maximal summation/subtraction (Herreras et al., 2023b).

Using numerical models (see Methods), we examined the spatial distributions of mixed FPs based on unidimensional spatial power FP profiles (as in ICA-derived FP-generators: volume conduction is implicit) normalized to equivalent strength and a realistic theta-frequency activity epoch (Figs. 5A, B). This qualitative analysis was limited to dipole polarity and spatial offsets and sought to identify which configurations are consistent with experimental observations of theta masking in the hilus. Relevant findings were as follows: (a) using spatial profiles from a realistic CA1 SLM generator and a DG generator, a “hilar” cancellation zone emerged only when these generators had opposing polarity configurations there (Fig. 5A, left and right panels); (b) spatial segregation of two different dynamics is best reproduced by adding a third, independent DG generator (Fig. 5B). It may also arise when one generator contained both theta and non-theta activity (e.g., gamma or beta) with opposing hilar polarity; however, in this case the non-theta activity inherits the spatial spread of theta and may propagate to CA1 or the thalamus, which is not observed experimentally (e.g., beta is not coherent between DG and CA1). Even if the LM and LPP gamma can spread to each others region, the rare phase-locking between these regional gamma activities would prevent efficient masking in the hilar region.

**Figure 5.**
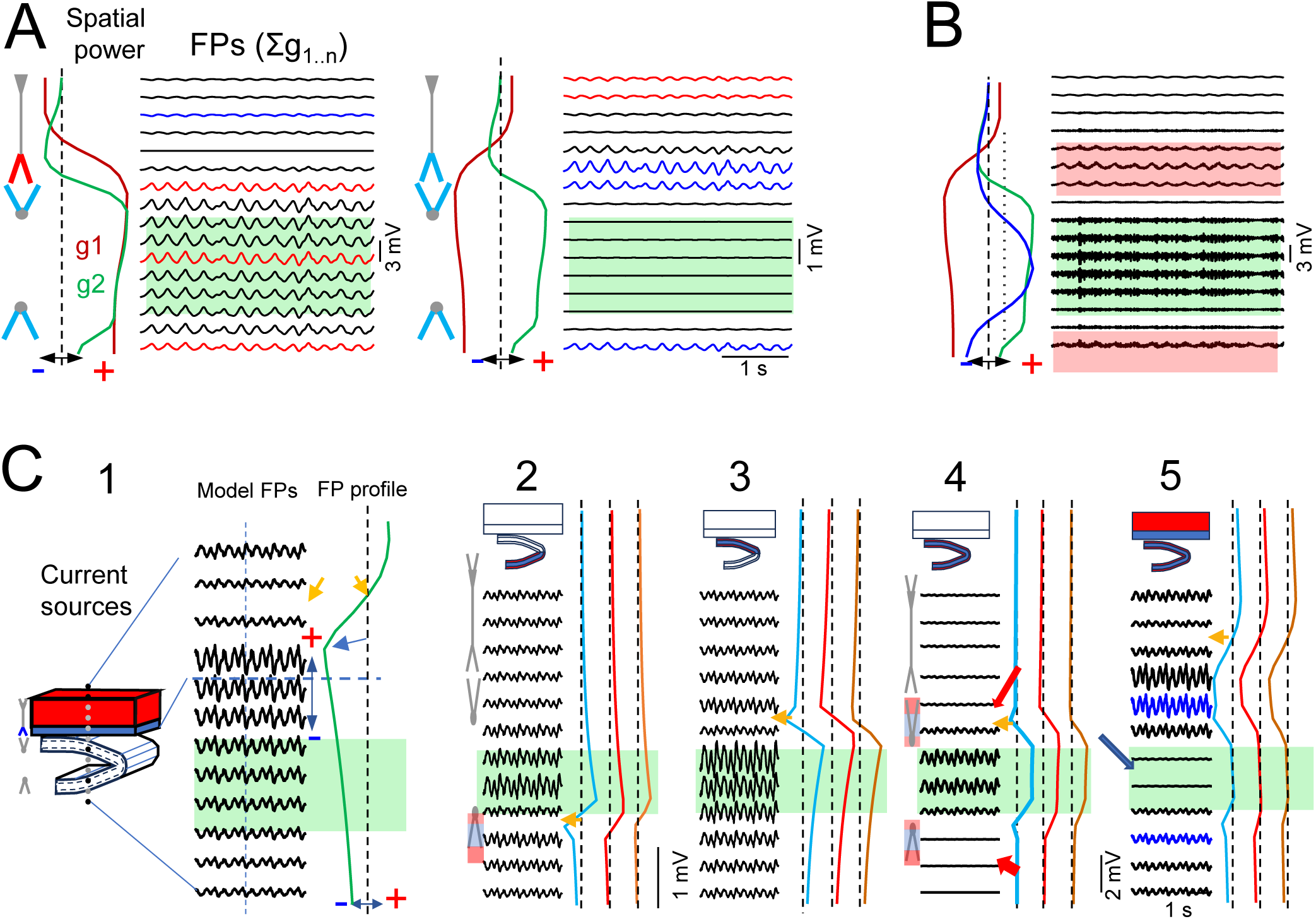
Boosting and cancellation in multisource FPs involving curved structures. Numerical (A,B) and FEM simulations (C). Linear FP profiles were generated by summing spatially weighted activities derived from realistic power distributions (A,B) or current source distributions based on experimentally characterized generators. A, g1 and g2 correspond to LM and DG generators, respectively, shown with either inverted (left) or experimentally derived polarities (right). Both generators shared the same theta-like temporal profile. Cell schematics indicate the active domains of pyramidal and granule cell layers and the polarity of the synaptic currents underlying each profile. FP cancellation (masking) occurred only at locations where opposite polarities and comparable amplitudes overlapped, requiring matching polarity at the distal dendritic domains of the original generators. Representative colored traces illustrate signal polarity at different recording sites. B, Spatial segregation of oscillatory components was optimal by introducing a third generator (blue profile) carrying distinct gamma (or beta) activity. C, Schematic blocks are colored when activated. The sites of polarity reversal are shown by orange arrows. Spatial current configurations correspond to the excitatory MPP (cyan) and LPP (red) inputs, and to inhibitory GCsom (brown) inhibition. (1) CA1 SLM-like excitation with a theta-like current pattern generates FPs that peak in the SLM and decay volume-conducted toward the DG (FP profile). These FPs are prominent across all DG regions, including the hilus and granule cell domains. Regional activation of either the upper (2) or lower (3) DG blades produces dipolar FP profiles. (4) Coherent theta-like activation of both DG blades produces large FPs in the interposed hilar region; note small FPs in granule cell source domains themselves (red arrows). (5) Only upon co-activation with the SLM generator do hilar potentials get masked.

To test these configurations under realistic volume conduction constraints, we used spatially defined synaptic sources within a volume conductor based on previously implemented finite element method (FEM) models of CA1 and DG (Torres et al., 2019), focusing on pathway-specific extracellular current geometry. Sources were modeled as quadrupolar (MPP) or dipolar (all others) according to their somatodendritic distribution (see Methods), and FPs were estimated along a linear track mimicking experimental recording (Fig. 5C). Current densities were assigned to 3D spatial blocks with pathway-specific geometries (Fernández-Ruiz et al., 2013) and intensity proportional to time fluctuations over realistic theta epochs with appropriate offsets to preserve polarity of currents across specific cell domains. Calculated FPs were AC-filtered and relative spatial power was estimated after ICA of linear FP samples.

Main results relevant for interpreting experimental findings follow: (a) Distal apical CA1 activation generated a dipolar field peaking at the CA1 border and passively spreading into the DG maintaining polarity (Fig. 5C,1); after AC filtering excitatory or inhibitory inputs yielded indistinguishable FPs, and when combined, fine-tuned phase alternation was required to minimize local cancellation; (b) single-blade GC activation produced hilar FPs with dipolar fields decaying smoothly on both sides (Fig. 5C,2-3), similar to activation in planar structures; (c) hilar-confined FPs (e.g., experimental beta or imaginary DG theta alone: Fig. 5C,4) required coherent input to both granule cell layers, whereas FPs in the granule cell source domains themselves is comparatively small (red arrows). This configuration produces minimal spread outside this region. (d) combining CA1 SLM input with any of the major DG pathways produced hilar masking when DG somatic currents (either passive or active) had opposite polarity to CA1 distal dendrites (Fig. 5C,5). In these configurations activity remains at both sides of the CA1/DG border and the lower blade, there reproducing experimental observations. No polarity reversal occurs across the DG, indicating that all DG theta is volume-conducted from CA1 whereas local DG theta is fully canceled (blue arrow). Whereas the excitatory LPP and the inhibitory GCsom (similarly oriented dipoles) were highly effective to cancel the volume-conducted CA1 theta in the hilus as they both drive large positive fields there, the quadrupolar MPP input required fine adjustment of laminar densities (asymmetric quadrupole). Combinations of co-activated MPP and GCsom inputs were also found effective with careful adjustment of relative laminar densities and power.

### The degree of FP cancellation depends on phase relationships between generators

Finally, we used a numerical approach to examine the impact of out-of-phase relationships between theta sources, as inferred from the transition from irregular to theta states (Fig. 1B). Theta epochs were generated by progressively increasing the phase lag (up to 180° of a cycle) and combined pairwise using the spatial power profiles of the CA1 LM and DG MPP generators (Fig. 6A). The mean FP amplitude was estimated at a location equivalent to the hilar region, ranging from 0.015 to 0.08 mV across a 180° lag arc (∼20% of the fully coherent amplitude: Fig. 6B). These simulations demonstrate that small differences in phase relationships between distributed generators can determine whether extracellular fields are amplified or cancelled. Notably, these estimates apply only to AC-coupled theta recordings and become less relevant for DC recordings when theta includes a sustained component (Brankack et al., 1993). Temporal fluctuations can also reduce power in the theta band as signals become more complex and exhibit higher frequencies in a space-dependent manner (Herreras et al., 2023b).

**Figure 6.**
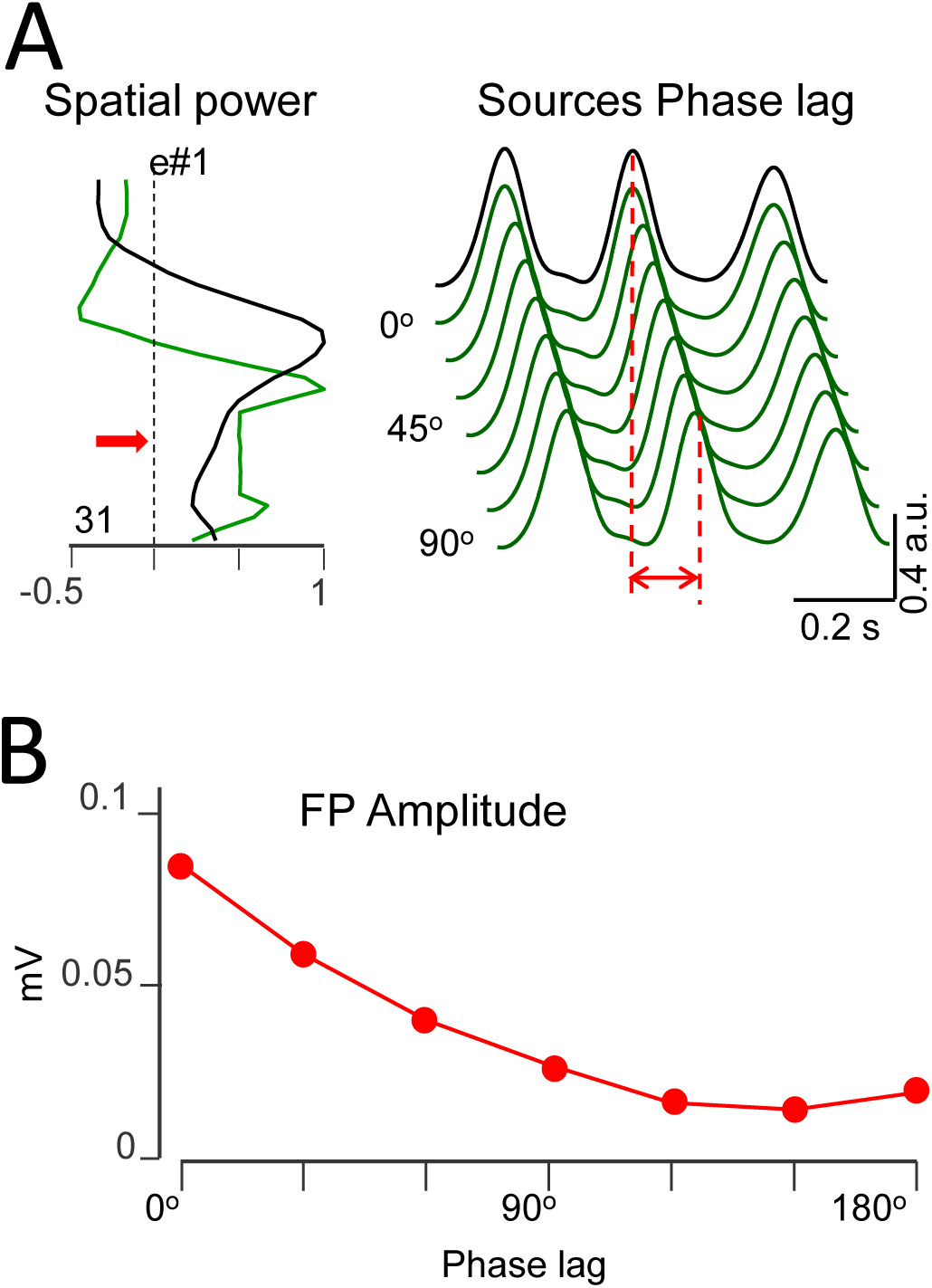
Phase lag between two theta sources produces variable cancellation in the summed FPs. A, the left panel shows the spatial power profiles. The red arrow indicates the position (hilar-like) where mean amplitude was calculated. The right panel shows a series of theta activations with increasing phase lag used as the second source in the pair. B, mean FP amplitude as a function of phase lag between the two sources.

## DISCUSSION

Here we show that the regional expression of hippocampal oscillations is determined by the interaction between anatomical geometry and the temporal coordination of distributed generators. Although theta generators are present in both CA1 and DG, their extracellular expression in the DG hilus is suppressed through geometry-dependent cancellation between coherent sources. In contrast, beta and gamma activities remain detectable because the corresponding generators lack stable interregional coherence and therefore do not undergo sustained cancellation. Together with computational modeling, these findings show that FPs reflect not only where neuronal currents are generated, but also how their geometry and temporal relationships shape their extracellular representation.

### Multiple theta sources, not all seen

A central objective of the present study is to emphasize that extracellular recordings provide access only to the subset of neuronal generators that become electrically expressed in the recording volume (Lorente de Nó, 1947). Understanding the mechanisms that suppress or distort extracellular fields is therefore as important as understanding those that generate them. From this perspective, volume conduction is to be regarded as the physical substrate that both enables and limits the emergence of mesoscopic extracellular fields. Theta-related activity provides an excellent framework in which these principles can be examined.

Theta oscillations are generally accepted to arise from multiple partially coherent generators distributed across the hippocampal–entorhinal system rather than from a unitary oscillatory process (Winson, 1974; Leung, 1984b; Montgomery et al., 2009; Lopez-Madrona and Canals, 2021). Present findings show that detectability of these generators depends critically on anatomical geometry and source interference, a phenomenon also observed in human cortex (Zuure et al., 2020).

The pathway specificity of mesoscopic FP generators implies that a single cell type may produce multiple generators (Herreras et al., 2015; Benito et al., 2014), as proposed for CA1 theta profiles (Holsheimer et al., 1982; Leung, 1984a; Balakrishnan and Pearce, 2014). In vitro and modeling approaches generate theta-like activity with spatial profiles that differ from in vivo depth recordings (Goutagny et al., 2009; Romani et al., 2024), which also vary across experimental conditions (Bland and Whishaw, 1976). Together, these observations indicate that theta field profiles depend not only on the underlying cellular mechanisms but also on how the corresponding generators combine in space.

This distinction between cellular activity and extracellular expression is often overlooked because generators are commonly inferred from either spiking activity or local field recordings, despite neither providing a direct description of the underlying current sources. A clear example is provided by the lateral habenula, where theta-modulated neurons fail to generate measurable local theta fields because their multipolar morphology leads to extensive cancellation of extracellular currents (Bertone-Cueto et al., 2020). Nevertheless, robust theta FPs are recorded in this region owing to volume conduction from the adjacent hippocampus. Indeed, theta-modulated neurons have been reported throughout the limbic system, including the nucleus accumbens, septum, amygdala, hypothalamus, thalamus and prefrontal cortex (Nuñez and Buño, 2021). Yet only a subset of these regions exhibit prominent theta FPs, and in several cases these have been proposed to reflect, at least in part, volume-conducted activity from hippocampal sources (Lalla et al., 2016; Roy et al., 2017; Torres et al., 2019; Kennedy et al., 2023). Thus, local neuronal activity and local FPs can become dissociated in both directions: active neurons may generate no measurable FP, whereas prominent FPs may arise from distant generators.

The hilus stands out as a notable, and somewhat paradoxical, exception, as it is known to exhibit large FPs when currents are coherently activated across both blades (Fernández-Ruiz et al., 2013). Although we show that theta currents are present and coherent in both blades, as for beta, gamma and other patterns, yet only theta failed to produce the expected large hilar potentials. Theta obscuring by concomitant large-amplitude beta oscillations is insufficient as hilar theta remains absent after beta-band filtering. Nevertheless, the lack of theta expression is not complete as reduced-amplitude hilar theta is observed during transient states in which the local power envelope varies independently of that at the main source in CA1 SLM. In addition, phase shifts between individual theta waves across recording sites become apparent during weak-theta states. Also, though not frequently mentioned, hilar theta can be observed in the literature in freely-moving animals (Buzsaki, 2002) suggesting the existence of multiple, partially independent theta generators.

### Observable DG theta largely reflects volume-conducted CA1 fields

Consistent with previous reports (Kocsis et al., 1999), pharmacological or functional blockade of entorhinal inputs reduces theta-related CSD activity in CA1 str. rad. and the DG molecular layer, while altering theta expression in CA1, supporting the existence of two major hippocampal theta generators located across the hippocampal fissure (Heynen and Bilkey, 1994; Ormond and McNaughton, 2015; Schlesiger et al., 2015; Zutshi et al., 2022). These findings have traditionally been interpreted in terms of changes in local generator strength, without considering how superposition of extracellular fields may affect downstream regions such as the hilus.

Weakening of one of these, the entorhinal component, shifts the balance toward CA1-dominated fields, producing a unimodal spatial profile centered in SLM that decays smoothly across the hilus, where CA1 theta becomes clearly visible, as predicted by feedforward and volume-conduction models (Herreras, 2016; Herreras et al., 2023a). Some qualitative features of the masked DG generator can be inferred, including a positive hilar polarity counterbalancing the negative one from CA1, and local inflections in cellular domains. These features are broadly consistent with the ICA-derived MPP component, whose theta activity is coherent with DG theta (Dickson et al., 1994; Deshmukh et al., 2010) and therefore with SLM activity. Thus, much of the theta observed in the DG reflects CA1 fields, while the local DG component remains masked. Because the C-shaped geometry of the DG reverses the apparent polarity of locally generated fields, this implies that the underlying CA1 and DG generators actually share the same intrinsic polarity.

### Seen or unseen, multiple DG theta generators

Because theta is oscillatory, conventional AC-coupled recordings remove baseline activity, causing CSD analysis to split individual theta waves into spurious source–sink sequences at each site, thereby limiting pathway identification (Brankack et al., 1993; Martín-Vázquez et al., 2013). Beta source–sink pairs are sharply localized to the outer two-thirds of the molecular layer, whereas DG theta currents extend across all granule cell domains. Accordingly, beta currents closely match an LPP origin (Chapman et al., 1998; Lockmann et al., 2018), while theta currents are more compatible with the theoretical quadrupolar distribution of MPP inputs arising from MEC layer II cells (Alonso and García-Austt, 1987; Mizuseki et al., 2009; Fernandez-Ruiz et al., 2013).

However, they could also reflect co-activation of excitatory and inhibitory inputs across different strata, which according to the model may generate diverse spatial distributions. Changes in the reversal site of residual theta currents after entorhinal input blockade further support the existence of multiple DG theta generators. Because MPP activation recruits hilar and basket interneurons that are themselves theta-modulated (Hafting et al., 2005; Scharfman and Myers, 2013), DG theta currents may arise from excitatory and inhibitory theta-related processes (Pernía-Andrade and Jonas, 2014), which remain difficult to disentangle experimentally. Multiple DG dipoles may also explain the failure of entorhinal input blockade to fully remove theta currents in this structure (Kocsis et al., 1999).

### The topology of activation makes geometry functionally consequential

Our results show that geometry alone does not determine extracellular field expression. Instead, geometry becomes functionally relevant only when distributed generators are sufficiently coordinated in time. Thus, multiple generators do not simply produce additive effects, but can lead to region-specific amplification or cancellation depending on geometry and coherence. Intermediate cases likely reflect a combination of structural and dynamical factors, including pathway topology and the spatiotemporal organization of synaptic inputs in a state-dependent manner (Benito et al., 2014). Thus, if FP amplitude depends mainly on the fraction of cells activated within a fixed pathway topology, mesoscopic source geometry behaves as an invariant and acts as a near-constant scaling factor. However, if activation is regionalized, effective geometry becomes state-dependent, strongly influencing both amplitude and phase of local recordings (Herreras et al., 2023a). Consistent with previous evidence of spatially heterogeneous recruitment in hippocampal networks, neuronal firing and synaptic inputs often display patchy and layer-specific activation rather than uniform engagement (e.g., place cells: Knierim et al., 2006; Mizuseki et al., 2009; Leutgeb et al., 2007). In such conditions, anatomy and geometry are not merely structural constraints, but active determinants of how oscillatory activity is expressed across regions.

### Generator location and field expression can become spatially dissociated

A remarkable finding is that the spatial expression of field oscillations does not necessarily identify the neuronal populations responsible for their generation. Thus, DG theta recorded in granule cell domains largely reflects CA1-generated fields, whereas hilar beta mainly originates from granule cell currents rather than hilar neurons. Such counterintuitive outcome emerges from the spatial geometry of the generators. As shown, both theta- and beta-related DG currents arise from synaptic inputs onto granule cells, yet beta field activity remains spatially confined to the hilus while theta is largely absent there. This creates a paradoxical dissociation between current generation and field potential expression, in which locally generated currents may remain electrically masked whereas propagated fields dominate the extracellular signal.

### Concluding remark

Together, these findings highlight that FPs are jointly shaped by neuronal dynamics, temporal coordination among distributed generators, and the geometry of the volume conductor through which those generators interact. Consequently, extracellular recordings do not necessarily reveal the location of the underlying generators, but rather their geometry-dependent extracellular representation. Accordingly, extracellular recordings provide reliable evidence for generators that become electrically expressed, but only limited evidence against the existence of generators that remain undetected. In fact, the absence of a field is itself a physiological observation whose explanation may reveal mechanisms that would otherwise remain hidden.

## Methods

### Experimental procedures

The experiments were performed in accordance with EU directive (2010/63/EU), and Spanish (RD 53/2013) and regional (Autonomous Community of Madrid, Order 4/8/1988) guidelines regarding the use of laboratory animals, and the experimental protocols were approved by the Research Committee of the Cajal Institute (I.D. Proex 367.5/21). Female Wistar rats (250–300 g, 3–4 months of age) were obtained from Janvier Labs (France). No evidence currently indicates that the biophysical mechanisms addressed here are sexually dimorphic. Yet, the conclusions are limited to the sex studied. Animals were housed in groups of four and provided with environmental enrichment objects within the cage. Rats were maintained under a 12 h light/dark cycle at controlled temperature (20–22 °C) with food and water available *ad libitum*. A total of 18 animals were used in two experimental sets, of which five were excluded from analysis due to inadequate electrographic state for present purposes. An additional group of 7 animals was reused from previous experimental series that underwent identical procedures (Hernández-Recio et al., 2023).

Animals were anesthetized with urethane (1.2-1.5 g/kg, i.p.) and placed in a stereotaxic device, and they were maintained hydrated by administering a glucose-saline solution (4 ml/kg) every other hour. The long-lasting anesthetic bupivacaine (0.75%) was applied to the surgical wounds. In different experiments concentric stimulating electrodes were placed in one of several areas to enable the recorded evoked potentials to be assigned to spatial landmarks: the soma layer of the CA3b region to activate the Schaffer input to the CA1; the olfactory tract to (di-synaptically) activate the LPP input to the DG; or the AB to activate the MPP input to the DG. Evoked potentials were also used to prime specific pathways and help identify FP generators (Korovaichuk et al., 2010; Hernández-Recio et al., 2023). Linear silicon probes (32 sites and 65 or 100 μm intersite distance: Atlas Neuroengineering, Leuven, Belgium) were placed at homotopic sites of both hippocampi across the dorsal hippocampus spanning the CA1 and DG/hilus regions (AP: -4.5; L: ±2.6-3). The probes were soaked in DiI before insertion (Molecular Probes, Invitrogen, Carlsbad, CA) to assess their location post-mortem in histological sections. A silver chloride wire implanted under the skin of the neck served as a reference for recordings. Signals were amplified and acquired using a MultiChannel System (Reutlingen, Germany) at a sampling rate of 20 kHz. At the end of each experiment, animals were euthanized with an overdose of urethane. Selected animals were then perfused through the abdominal aorta with PBS followed by 4% paraformaldehyde. Sagittal brain sections (100 µm) were stained with bis-benzimide to assess electrode positions by fluorescence microscopy. In an experimental series we aimed at selectively blocking the entorhinal input to the hippocampus in one hemisphere by delivering 1 μl of the anesthetic lidocaine (HCl 2%: Braun) at the AB using the outer cannula of the stimulating electrode. This was connected to a 5 μl Hamilton syringe through a P20 PE-tubing.

Due to fanning out of perforant path fibers, single injections may not completely disrupt the entorhino-hippocampal communication. Thus, we used repeated 0.2 μl ejections every 200 μm in an ascending trajectory to minimize loss of solution due to backflow. Effective disruption was considered when evoked fEPSPs in the DG were abolished and the spontaneous activity of the LPP and MPP generators —separated off-line as indicated below— in the ipsilateral hippocampus could no longer be observed for at least 15 min, but remained in the contralateral one. These conditions were met in five out of seven animals.

### Signal treatment and analysis

#### Signal selection

The selected signals were wideband FPs (0.1 Hz - 5 kHz), and they were resampled at 10 kHz. The achieved anesthetic plane permitted a wide variety of short-lived electrographic irregular patterns interspersed with epochs of spontaneous theta. In such conditions, theta could be readily elicited by gentle prick of the animal’s tail. No difference was found between spontaneous and evoked theta and they were treated similarly. For reference we defined frequency bands as follows: delta (0.5-2 Hz), theta (3-5 Hz); alpha (9-11 Hz), beta (15-20 Hz), and gamma (30-50 and 60-90 Hz: low and high, respectively). Theta epochs were selected when the theta band of the period considered reached a rhythmicity index of at least 0.7, calculated through the decline of the first peaks in the autocorrelation function (ACF) as follows:

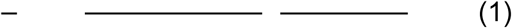

where A(n) is the peak of the ACF (N = 5). RI is bounded so that RI = 1 corresponds to sine waves, whereas 0 is for noise. For a reference, persistent theta in the SLM of CA1 has RI ̴ 0.8.

#### Spatial profiles of different activities

Selected epochs of FPs may exhibit characteristic depth profiles across the hippocampus, as in evoked potentials. In laminar structures, spatial profiles of FP activity provide a rough approximation of source location, although they may be strongly shifted in curved populations (Woodbury, 1960; Herreras et al., 2023a, b). To construct profiles, the mean power (V²) over a 100 s period at each recording site. Wideband and band-pass (theta, beta, and gamma) filtered FP profiles were built, and they were aligned across individuals with respect to an easily identifiable site in the SLM where theta exhibited maximal power. No further spatial normalization was performed, given the variability in interlaminar distances across recording tracks and animals. Evoked potential profiles were constructed at the peak of the fEPSP following MPP activation (averages of n = 10 stimuli). Care was taken to reduce spike contamination by using a stimulus intensity 50% of the spike threshold. In addition, spatial profiles for pathway-specific FP generators were obtained using ICA (Herreras et al., 2015).

#### Separation of mixed sources by independent component analysis

Since the FPs produced by synaptic inputs to the CA1 SLM spread into the DG hilus and overlap there with entorhinal driven granule cell generated FPs they add up variably at different sites according to spatial power and polarity. Thus, we required the regional activities to be separated in order to confirm the selective chemical blockade of only one set, as both carry theta oscillations in our FP samples of interest. This was achieved through the ICA as in former studies (Makarov et al. 2010; Benito et al., 2014). In the hippocampus, ICA greatly facilitates the resolution of the different voltage gradients produced by distinct synaptic inputs even if they reach the same cell population. Most hippocampal FP-generators are not only population-specific, but also pathway-specific (Herreras et al., 2015).

We employed the kernel density ICA algorithm (KDICA) (Chen, 2006) customarily implemented in MATLAB (the package is available at: http://blogs.mat.ucm.es/vmakarov/downloads). Recorded FP signals *u_m_(t)* were considered as the weighted sum of the activities of *N* neuronal sources or FP-generators:

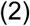

where (*V_mn_*) is the mixing matrix composed of the so-called voltage loadings -or spatial weights- of *N* FP-generators on *M* electrodes, and *s_n_(t)* is the time course of the *n-th* FP-generator. Thus, the raw FP observed at the *m–th* electrode is a linear mixture of the electrical activity of several independent FP-generators. Using {u_m_(t)}, the ICA finds both the matrix (V_mn_) and s_n_(t). The joint group of spatial weights (V_mn_) is ordered into instant depth profiles of the voltage according to electrode position and reflects a linear sample of the 3D voltage shell generated by a given source. Such profiles can be compared to the spatial profiles of the standard evoked potentials of specific pathways when available. In fact, both spontaneous and evoked potentials are captured in the temporal envelope of an FP generator (Korovaichuk et al., 2010; Benito et al., 2014).

Although a single linear profile is insufficient to infer the full three-dimensional voltage field, it provides critical information on gradients in a given direction that reflect whether electrode arrays are within or close to the source (accelerating curved spatial gradients) or far out (non-zero flat profile) (Herreras et al., 2023a). The time-course of an FP-generator *s_n_(t)* can be considered as a postsynaptic temporal convolution of spike output in an afferent population (i.e. afferent spike trains), as shown experimentally for the CA3-CA1 pathway (Fernández-Ruiz et al., 2012) and numerically for a variety of inputs (Makarova et al. 2011; Martín-Vázquez et al., 2013, 2016). The mathematical validation, robustness and practical limitations of this approach, as well as the possible causes leading to faulty separation and some of the possible workarounds have been thoroughly investigated using feed-forward models of FPs with realistic geometries (Makarova et al., 2011; Martín-Vázquez et al., 2016; Torres et al., 2019). A compound variance > 99% of the original FP variance was always retained.

#### Current source density analysis

Current source density analysis is commonly used to reduce the contribution of volume-conducted potentials originating outside the recording region (Lorente de Nó, 1947; Leung, 1979). Here, we employed CSD primarily to assess the presence or absence of local transmembrane currents at specific recording sites. Assuming constant extracellular conductivity (σ), CSD was estimated as *CSD = –σ ^2^u*, where *u(t,x,y,z)* is the extracellular potential and is the Laplacian operator. For linear probes (typically 32 channels), we used a one-dimensional approximation based on the second spatial derivative of the voltage profile along the main cell axis.

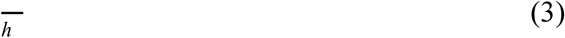

where *u_m_(t)* is the FP recorded at the *m-th* site and *h* is the inter-site distance.

In contrast to evoked responses, where a pre-stimulus baseline can be defined, CSD estimates from spontaneous FPs recorded in AC-coupled mode lack a true reference level. As a result, the polarity, phase, and amplitude of CSD signals are not reliable for quantitative analysis (Brankačk et al., 1993; Martín-Vázquez et al., 2013). Moreover, CSD traces at a given site still reflect mixtures of sinks and sources arising from multiple pathways with overlapping spatial domains and independent dynamics, limiting their interpretability (Hernández-Recio et al., 2023).

### Forward model of field potentials

#### Finite Element Models

Layer curvature has a marked effect on the spatial distribution of extracellular fields compared to planar structures (Woodbury, 1960; Herreras et al., 2023a), distorting the field distribution throughout the surrounding volume and leading to non-intuitive alterations in laminar FP profiles. Former theoretical models for exploration of theta profiles were far too simple and did not consider layer curvatures. Thus, we used forward modeling of FPs in the CA1 and DG regions using standard and fake geometries to better understand the 3D spatial expansion of their theta FP generators and the mutual distortions during co-activation (Torres et al., 2019; Muñoz-Arnaiz et al., 2025). It is worth noting that FP modeling does not require the explicit anatomy of cells and populations, but rather the current sources they generate. In layered populations, such as the CA1 pyramidal and granule cells, pathway-specific extracellular current sources can be well reproduced by FEM models, which allow explicit modeling of the geometry and the dielectric properties of the extracellular space. We modified a formerly deployed model of FPs generated by the main hippocampal regions in a rat brain hemisphere built with COMSOL Multiphysics software (COMSOL, Inc., Burlington, MA) (Torres et al., 2019).

Sinks and sources of extracellular current (Lorente de Nó, 1947) were built as rectangular slabs of homogeneous current density spatially assembled to match the population curvatures. We used the physical dimensions of the different modeled structures from the rat brain atlas (AP 4.56mm: Paxinos and Watson, 2007). Two sets of blocks were used for theta generators, mimicking the LM-to-CA1 and MPP-to-DG synaptic inputs, respectively. Additional inputs to the DG (LPP and a somatic inhibition) were also built and combined discretionarily. The blocks of current were enclosed in a surrounding volume of “extracellular space” of 20 × 20 × 20 mm (H, W, L), which minimizes the distortion of field curves in the region of interest (Fernández-Ruiz et al. 2013). For simplicity, the curvatures of the DG were implemented in only one direction that served well the main objective. We used Dirichlet boundary conditions on the surface enclosing the volume and imposed charge conservation inside. A tetrahedral adaptive mesh of the highest resolution (smallest size: 30 μm) was used to ensure the correct resolution of field equations in the curved compartments.

The laminar distribution of currents for each source was approached by three or four stacked slabs whose amplitude and density were obtained experimentally by CSD analysis of evoked potential profiles or the FP generator profiles along the main axis of the respective principal cells (Benito et al., 2014; Torres et al., 2019). The SLM, LPP, and GCsom inputs were configured as dipolar (excitatory or inhibitory inputs to a distal domain of the principal cell), whereas the MPP was configured as quadrupolar (active sink domain in the middle third of the GC dendritic tree). The current density of source and sink stacked slabs was balanced to reflect zero net current across the domains of single cells, and it was manually adjusted to achieve 0.6-1 mV theta waves in the SLM or 60-120 µV irregular series of waves in the center of the active sink layer (Chauvette et al., 2010; Torres et al., 2019).

The dynamics for each generator were obtained from actual experiments and epochs. The inhibitory generator in the DG, with an active source layer facing the hilar side, was composed of irregular activity with interspersed 100-200 ms-long waves and superimposed bursts of alpha-beta waves, jointly reaching 0.8-1 mV in the hilus (Benito et al., 2014).

Model FPs were estimated in vertical tracks with a spatial resolution of 50 μm in dorso-ventral tracts. Non-realistic profiles in simulated tracks were also employed to challenge ML techniques on different blends of FPs elicited by the same generators as well as to test the CSD and ICA performance on synthetic and more naturalistic signals.

#### Numerical estimation of FPs

Assuming quasi-static conditions and homogeneous isotropic conductivity in volume conductors, extracellular FPs can be described as a linear superposition of contributions from multiple current sources (Eq. 2). Within this framework, we implemented a simplified forward model operating directly in the space of experimentally recorded signals. For each synaptic pathway, we used laminar FP profiles obtained experimentally via ICA, which represent the effective contribution of each pathway after volume conduction. These profiles are treated as stationary in time, reflecting a stable underlying distribution of transmembrane currents under given experimental conditions.

Simulated signals were generated by constructing laminar matrices for each pathway and performing channel-wise linear summation to obtain mixed FPs. Individual profiles were normalized to their maximum amplitude prior to summation to facilitate comparison across configurations. Temporal activation patterns were derived from short experimental epochs of theta activity, and beta and gamma dynamics were also incorporated either independently or in combination with theta. The main parameters explored were relative polarity, amplitude, and phase relationships between contributing pathways. Simulations were performed along linear recording tracks with a spatial resolution of 50 μm. The goal of these simulations was not to reproduce absolute field amplitudes, but to identify the conditions under which linear interference between pathway-specific generators leads to partial or complete masking of theta activity.

### Statistics

Mean and standard error of the mean were used for all quantifications. Statistical comparisons used the two-tail t-test and significance was established at 0.95 %. For group comparisons we used ANOVA and post-hoc Bonferroni corrections, with Fisher z-transformation for group significance.

The time evolution of the power of an FP-generator (in mV^2^) was calculated by:

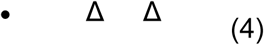

where *v*(*t*) is the virtual FP at the electrode with maximal power and *Δ* is the length of averaging (60 s).

For visualization of frequency content, we estimated the periodogram power spectral density of the temporal activation of FPs or FP-generators (power spectra) and then computed the signal power in different frequency bands (in Hz) as indicated above.

Coarse synchronization between different recording sites and time courses of FP-generators was estimated using the cross-correlation coefficient (CC) and spectral coherence. The CC index was obtained as:

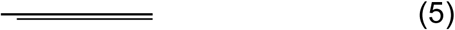

The spectral coherence of the activity of two FPs or CSD time traces was calculated by

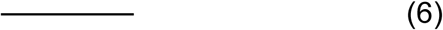

where {*P_xy_(f)*} is the matrix of cross-power spectral density. To determine the level of significance we used the surrogate data test. Randomizing phase relations and keeping other first order characteristics intact, we obtained surrogate time series from the original signals. For each experiment we generated 1,000 surrogates and evaluated spectral coherence. The significance level (at α = 0.05) was then calculated for each frequency value and coherence above this level was considered statistically significant.

We also evaluated the probability distribution histograms (PDH) of the phase difference between time activations of two FPs or CSD time traces *Δϕ(t) = ϕ_2_(t) - ϕ_1_(t)*. For proper estimation of the activation phase *ϕ_n_(t)* we used down sampling of *s_n_*(*t*) followed by band-pass filtering in the range 2-10 Hz or 10-20 Hz for theta and beta bands, respectively. The signal obtained was passed through the Hilbert transform, yielding the phase *Δϕ_n_(t).* Using *Δϕ(t)* we calculated the PLV between FPs or CSD traces:

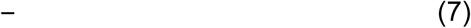

The PLV attains a value of 1 if, and only if, the signals are phase-locked (synchronized), whereas PLV → 0 if the signals are uncorrelated. To examine the statistical significance of the PDH and PLV, we adopted a surrogate data test. On this purpose, time series were obtained from the original signals by randomizing the phase relationships and keeping the other first order characteristics intact, identifying a 95% confidence interval from the surrogate distribution. For PDH we used the Bonferroni correction to avoid the growth of false positives. If the PDH or PLV value exceeded the critical confidence limit then we decided positively on the presence of phase coupling between FPs. In this case, the maximum PDH was also used to evaluate the phase lag between two signals (*Τmax*).

Spatial maps of linear correlation were built by estimating the Pearson correlation pairwise through all recording channels. Values fluctuate in the interval [-1, 1], where absolute values close to 1 indicate a good correlation while values near 0 indicate that the variables have no correlation at all. Pseudocolor maps were built for ease of regional correlation, aided by anatomical and electrophysiological landmarks.

## Author contributions

JM and FA performed the experiments; JM and RMA performed model studies; OH conceived the study and wrote the original draft; all authors analyzed the data, reviewed and approved the manuscript.

## Data availability

The datasets generated and/or analyzed during the current study, together with the source code required to reproduce the analyses, are available from the corresponding author upon reasonable request.

## Supplementary Material

**Supplementary Figure 1.**
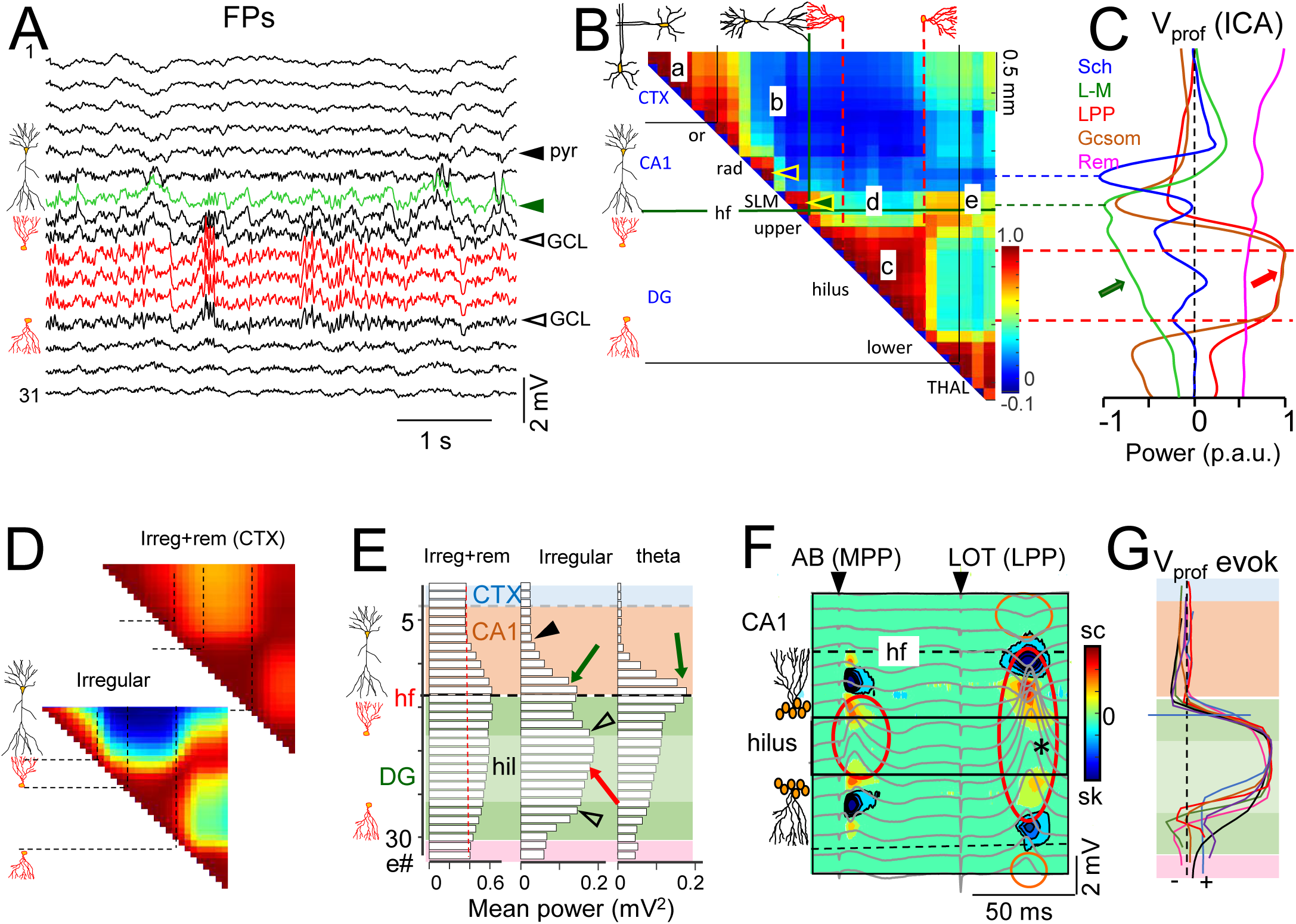
Spatial features of hippocampal FPs during non-theta states. A, Sample traces of linear FPs across CA1–DG. Hilar FPs (red) show similar dynamics. The green trace corresponds to the CA1 SLM. One of every other trace is shown (n = 32). Cell drawings in the margins indicate anatomical layer landmarks. B, Spatial map of pairwise cross-correlations. Zones of high coherence (a,c) are typically local, but also occur between separated sites (zone d). C, Normalized spatial profiles of ICA-disentangled FP generators (color-coded), with anatomical landmarks and correspondence to CC maps. Arrows indicate decaying hilar activity for the LM generator and steady activity for the DG generator. D, CC maps corresponding to two patterns of irregular hippocampal activity, with (upper) and without (lower) cortical slow-wave activity. The high, uniform correlation indicates dominance of volume-conducted cortical activity across sites. E, Sample profiles of power across individual channels for three EEG states. The black arrowhead marks the CA1 stratum pyramidale (st. pyr). Open arrowheads mark granule cell layers in the DG. Green and red arrows indicate local peaks of power at the distal CA1 SLM and the hilus, respectively. F, Paired-pulse evoked potentials from the angular bundle (AB) and the lateral olfactory tract (LOT), superimposed on CSD sources (yellow/red) and sinks (blue). Note the larger hilar FPs (red oval) without corresponding CSD transients in the hilus (asterisk), and the volume conduction of potentials out of the DG (green ovals) only under dipolar current configurations. G, Superposition of AB-evoked potential profiles from seven animals, aligned at the 50% amplitude site in the upper DG blade. Abbreviations: CTX, cortex; THAL, thalamus; or, pyr, rad, SLM: oriens, pyramidal, radiatum, and stratum lacunosum-moleculare; GCL, granule cell layer; hf, hippocampal fissure; hil, hilus; Vprof, voltage profile; sc/sk, source/sink; p.a.u., proportional arbitrary units. Color codes and shaded zones are consistent across figures.

**Supplementary Figure 2.**
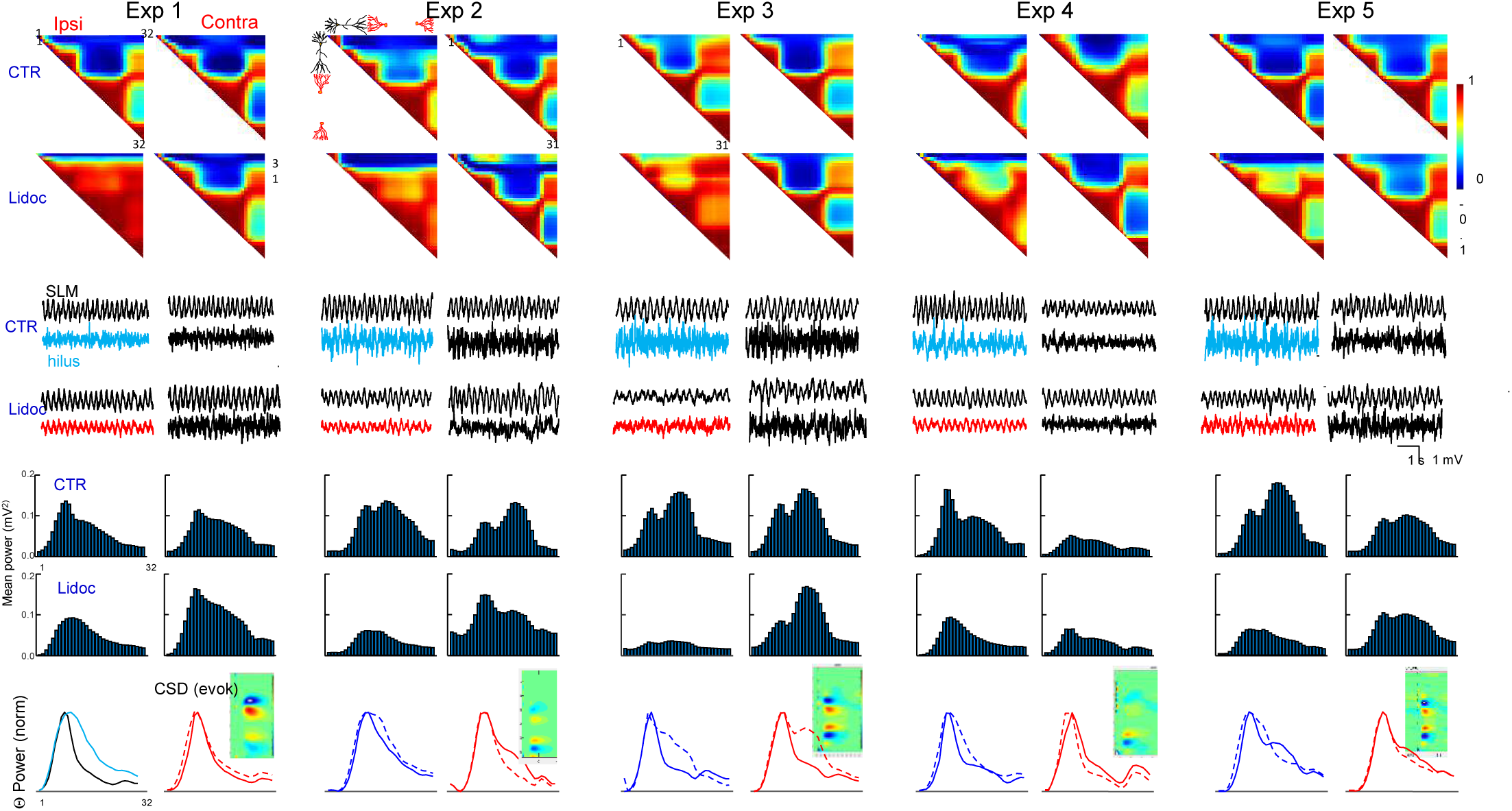
Unmasking of hilar theta by lidocaine injection into the angular bundle: additional experiments. Data processing was performed as described for Fig. 4 in the main text. For each experiment, two columns of panels are shown corresponding to recordings from the left (ipsilateral) and right hippocampus (contralateral to the lidocaine injection). Control conditions are displayed in the upper panels, whereas the effects of lidocaine are shown in the lower panels. From top to bottom: spatial maps of pairwise Pearson correlations; representative FP traces recorded in the SLM and hilus; mean power profiles; and normalized theta power profiles (solid and dashed lines correspond to control and lidocaine conditions, respectively). Insets show CSD maps corresponding to AB-evoked potentials in the DG before lidocaine injection.

